# Impact of early locus coeruleus lesions in the TgF344 Alzheimer’s disease rat model

**DOI:** 10.1101/2025.11.09.687363

**Authors:** Alexia E. Marriott, Jason P. Schroeder, Anu Korukonda, Brittany S. Pate, Katharine E. McCann, David Weinshenker, Michael A. Kelberman

**Author notes:** Address correspondence to: Michael A. Kelberman, Department of Physiology and Neuroscience, University of Colorado, Boulder, Gold Biosciences A220C, Colorado Ave, Boulder, CO 80309.

## Abstract

**INTRODUCTION:** In murine models of Alzheimer’s disease (AD), lesioning the locus coeruleus-norepinephrine (LC-NE) system with DSP-4 exacerbates AD-like neuropathology and cognitive impairment. However, the impact of LC lesions during prodromal stages is poorly characterized.

**METHODS:** TgF344-AD and wild-type rats received monthly injections of DSP-4 or saline from 1-5 months of age, a time point preceding forebrain plaque or tangle deposition in TgF344-AD rats, after which behavior and pathology were assessed.

**RESULTS:** DSP-4 compromised LC cell bodies, fibers, and NE content. LC lesion and the AD transgene each affected several affective behaviors and/or cognition individually, but few interactions were found and DSP-4 failed to exacerbate behavioral phenotypes or neuropathology in TgF344-AD rats.

**DISCUSSION:** Combined with previous literature, our data suggest that LC lesions exacerbate pre-existing AD-like pathology and behavioral impairments, rather than accelerate their onset. Further characterization of LC lesions in TgF344-AD rats at different ages is warranted.

**Research-in-Context:**

1. Systemic review: The authors reviewed existing literature using traditional sources, including PubMed. Previous studies investigated the impact of locus coeruleus (LC) lesions on Alzheimer’s disease (AD)-like neuropathology and behavior in murine models of AD. However, the impact of LC degeneration in an animal model that expresses both amyloid and endogenous tau pathology at a time point before the emergence of significant forebrain pathology is underexplored.
2. Interpretation: We expanded the behavioral and molecular characterization of TgF344-AD rats in response to N-(2-chloroethyl)-N-ethyl-2-bromobenzylamine hydrochloride (DSP-4)-induced LC lesions during the pre-pathology stages of disease. Unexpectedly, we found that TgF344-AD genotype and DSP-4 rarely interacted to exacerbate AD-related symptoms or pathology.
3. Future Directions: Our results indicate LC lesions do not accelerate onset of AD-like neuropathology or behavioral impairment in this model. Future studies in older TgF344-AD animals and using different DSP-4 treatment regimens would help clarify the relationship between LC integrity and AD progression.

**Highlights:**

- Locus coeruleus damage causes apathy-like behavior and changes in arousal
- TgF344-AD genotype induces social recognition deficits and anxiety-like behavior
- Locus coeruleus damage and TgF344-AD genotype rarely interact to worsen deficits
- Interactions that exacerbate Alzheimer’s disease neuropathology were also scarce

## 1. Introduction

The locus coeruleus (LC) is a bilateral nucleus in the dorsal pons and is the primary source of norepinephrine (NE) in the central nervous system. LC neurons, which influence an array of behaviors including arousal, stress responses, mood, and cognition, are uniquely susceptible to insults triggered by neurodegenerative disorders [1]. It has long been recognized that loss of noradrenergic neurons in the LC is a universal feature of Alzheimer’s disease (AD), and studies by multiple groups indicate that LC neurons are the first to accumulate hyperphosphorylated tau in the brain and show signs of fiber damage in the earliest stages of AD [2–4]. While LC neurons eventually succumb to pathology, frank cell loss only begins in mid-late stages of disease [5,6].

Spatiotemporal patterns of tau accumulation and neuronal loss indicate a substantial period of time in which LC dysfunction could contribute to aspects of AD, which has been the major focus of previous studies. For example, NE-sensitive symptoms such as anxiety, depression, and sleep disturbances are common in prodromal AD and are temporally linked to the appearance of aberrant tau in the LC [1,7,8]. Our lab previously reported age-dependent accumulation of hyperphosphorylated tau in the LC and changes to LC firing rates in TgF344-AD rats that overexpress mutant amyloid precursor protein and presenilin-1 [9,10]. Specifically, footshock-induced LC firing is elevated in young TgF344-AD rats at an age in which anxiety-like behavior emerges, while the LC is hypoactive in old TgF344-AD rats that show deficits in reversal learning. These age-dependent differences in firing rates support the idea that LC hyperactivity is linked to prodromal neuropsychiatric symptoms of AD, while deficient LC-NE transmission may contribute to cognitive impairment [1,8,9]. Further support comes from human literature, where early neuropsychiatric symptoms are associated with increased cerebrospinal fluid NE and metabolite levels, while decreased LC integrity predicts cognitive decline [11–13]. There are also positive correlations between the magnitude of neuronal loss in the LC and both AD duration and severity [3,14].

Based on this evidence, studies have used the selective LC neurotoxin N-(2-chloroethyl)-N-ethyl-2-bromobenzylamine hydrochloride (DSP-4) to isolate phenotypes related to LC dysfunction and degeneration in neurodegenerative disorders. These studies show that DSP-4 induces LC lesions that vary in severity based on specific treatment regimen [15–17], but in general lesions axons, can damage cell bodies, and produces neuroinflammation [18–21]. When DSP-4 has been applied in amyloid-and tau-based rodent AD models, LC lesions exacerbate AD-like neuropathology and cognitive impairment [15–17,20,22–24]. However, most studies use DSP-4 lesions in AD models at time points when pathology has already accrued. This leaves an open question as to whether early LC degeneration accelerates the onset of AD-associated behavioral impairment and pathology.

The TgF344-AD rat has unique features compared to other rodent AD models [25]. These rats display both amyloid and tau pathology that is age-dependent, including aggregation of endogenous hyperphosphorylated tau in the LC prior to pathology in other brain regions [10]. This mirrors a hallmark aspect of the human condition often absent in other rodent AD models. Crucially, pathological changes in the LC in human AD occur decades prior to cognitive decline associated with bona fide plaques and tangles in the forebrain [2]. Thus, there is potential for LC-based therapeutic interventions, provided that such changes are detected early enough. To achieve this, a more nuanced understanding of LC pathological changes and their impact on disease progression are needed, especially how LC degeneration in the absence of significant pathology may accelerate disease progression. We tested this by inducing LC-specific damage in TgF344-AD rats before the appearance of amyloid, tau, or neuroinflammation in the forebrain, and examined its impact on AD-related pathology and behavior.

## 2. Materials and Methods

### 2.1 Animals

Male and female TgF344-AD rats and their non-transgenic wild-type (WT) littermates were used in this study. The animals were housed in groups of 2-3 until behavioral testing, at which point they were single housed. Lights were on a 12-h cycle (lights on at 7:00 AM) and rats had *ad libitum* access to food and water, except during behavioral testing when specified. The Emory University Institutional Animal Care and Committee approved all the experimental procedures used in this study.

### 2.2 DSP-4 lesions

The current study consisted of four experimental groups: Saline-treated WT (n=11, 6 males, 5 females), DSP-4-treated WT (n=9, 5 males, 4 females), saline-treated TgF344-AD (n=12, 6 males, 6 females), and DSP-4-treated TgF344-AD animals (n=9, 2 males, 7 females). At ∼1 month of age, animals received two injections of saline or DSP-4 (50 mg/kg, i.p.) spaced 1 week apart. Animals then received saline or DSP-4 injections once per month for the next 4 months until they were ∼5 months old. For monthly booster injections, animals were lightly anesthetized using isoflurane. Behavioral testing began 1 week following the final injection of saline or DSP-4.

### 2.3 Behavioral Assays

#### 2.3.1 General

Behavioral assays were conducted to assess changes in social behavior (social discrimination paradigm), circadian activity (23-h locomotor activity), arousal (novelty-induced locomotion, sleep latency), stress-induced repetitive behavior and environmental engagement/apathy-like behavior (nestlet shredding, stick chewing), anxiety-like behavior (open field and novelty-suppressed feeding) and learning and memory (cue-and context-dependent fear conditioning), many of which have been described previously by our group [8,26]. Behavioral tests were separated by at least 1 day and performed in the following order, from least to most stressful, during the light phase: social behavior, locomotor activity, sleep latency, open field, novelty-suppressed feeding, and cue-and context-dependent fear conditioning. Stick chewing began during locomotor testing, while nestlet shredding started at the beginning of food deprivation for novelty-suppressed feeding.

#### 2.3.2 Social discrimination

Rats were individually placed into a 22.5” l × 17.5” w × 12” h rectangular, opaque plastic arena. A plastic divider and wire mesh were used to split the arena into 3 distinct chambers. Wire mesh allowed non-contact investigation but prohibited physical interaction within the social zones. For 2 days prior to testing, rats were habituated to the arena for 10 min per day. On test day, rats were habituated for 10 min in the arena. Rats were then removed for 5 min. Rats were placed back into the arena for 10 min, this time with two plastic rats on one side and a novel, same-sex, approximately age-matched WT stimulus animal on the opposite side behind the wire mesh walls. All animals were then removed for 5 min. Finally, rats were placed back in the chamber for 10 min. The familiar stimulus rat was placed in the same chamber as in the previous trial, and another novel, same sex, approximately age-matched, WT stimulus animal was placed in the other chamber behind the wire mesh walls. Chambers were cleaned with Vimoba between trials and side of novel animal/object was randomized and counterbalanced. Activity was videotaped and scored using ANY-Maze software (Version 7.47). The primary outcome measure was the duration of time spent in specific chambers of the arena. The following formula was used to calculate the ratio of time spent investigating the novel animal versus novel object:

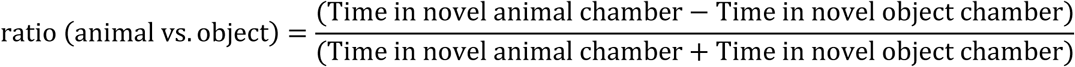

Similarly, the following formula was used to calculate the ratio of time spent investigating the novel animal versus the familiar animal:

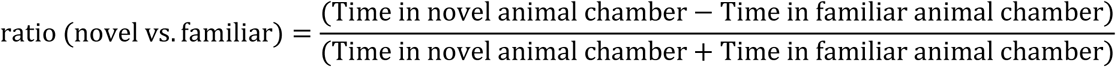

#### 2.3.3 Circadian locomotor activity

Rats were placed in a new cage with fresh food, water, and clean bedding. Cages were equipped with external infrared photobeams to measure locomotor activity. Total number of ambulations were binned in 15-min intervals over the 23-h testing period. The primary outcome measures were the number of ambulations during the light and dark phases and total number of ambulations over the entire 23-h period. In addition, the first 30 min of the circadian locomotor testing were considered novelty-induced locomotion as a general level of arousal.

#### 2.3.4 Stick chewing

At the start of circadian locomotor activity, a pre-weighed wooden stick was placed into the cage and was weighed after 24 h and 1 week. The amount of stick chewed was used as a proxy of apathy-like behavior.

#### 2.3.5 Sleep latency

Rats were removed from their home cage, gently handled, and placed back into their home cage which was covered with a transparent sheet of Plexiglass for overhead video recording. Animals were recorded for 4 h, and latency to fall asleep was assessed. Sleep was operationally defined as at least 75% sleep for 10 min, the first 2 min of which were uninterrupted. Sleep was determined by a combination of closed eyes, sleep posture, and immobility. These combined criteria are an EEG-verified measure of sleep in rodents [27,28].

#### 2.3.6 Open field

Rats were individually placed in the center of an open field arena (44” l × 33.5”w × 10.75”h) and videotaped for 5 min while they freely explored the arena. The amount of time spent in the center of the arena was used as a measure of anxiety-like behavior, with more time in the center interpreted as lower anxiety-like behavior.

#### 2.3.7 Novelty-suppressed feeding

24 h prior to testing, rats were food deprived in their home cage. At the time of testing, a single pre-weighed food pellet was placed on a piece of weigh paper in the center of a rectangular arena (22.5” l × 17.5” w × 12” h). Testing was performed in the dark with the arena lit under red light. Rats were placed in the arena, and the latency to eat in this novel environment was timed. The session concluded once a rat bit into the pellet or 25 min had elapsed, whichever occurred first. Immediately after, the rat was placed back in its home cage with the same pellet, and the latency to eat in this familiar environment was recorded. To control for hunger levels, the amount of pellet consumed in 1 h following the behavioral test was also measured.

#### 2.3.8 Nestlet shredding

At the beginning of food deprivation for novelty-suppressed feeding, a pre-weighed nestlet was placed into the home cage. The un-shredded portion of the nestlet was weighed after 90 min and 6 days. The amount of nestlet shredded in the first 90 min was used as a measure of stress-induced repetitive behavior, while the 6 day time point was used as another proxy of apathy-like behavior.

#### 2.3.9 Fear conditioning

Testing was performed over the course of 3 days, with training on the first day, contextual fear conditioning on the second day, and cued fear conditioning on the third day.

Training: On the first day, rats were placed in a fear conditioning chamber containing a floor composed of stainless-steel bars that was cleaned with Vimoba. Rats were allowed to habituate to the chamber for 3 min. Following this 3 min habituation period, an 85 dB tone sounded for 20 s which co-terminated with a 2 s, 0.5 mA footshock. This was repeated two more times, with a 1 min inter-trial interval in-between each footshock. Chambers were cleaned using Vimoba between sessions.

Contextual fear conditioning: On the second day, rats were placed in the same chamber used during training for 8 min without the presence of a tone or footshock. The amount of freezing was recorded. Chambers were cleaned using Vimoba before testing and between sessions.

Cued fear conditioning: On the third day, rats were placed in the fear conditioning chambers with a novel texture (metal grid flooring) and odor (chambers cleaned with 70% ethanol) and allowed to habituate for 2 min. Following this 2 min habituation period, an 85 dB tone was played for 20 s in the absence of footshock. This was repeated two more times, with a 1 min inter-trial interval between each round. The amount of freezing was recorded.

### 2.4 Tissue Preparation

Animals were sacrificed following behavioral testing, being randomly assigned to a tissue collection method while attempting to balance by sex within a specific experimental group. All animals were deeply anesthetized using isoflurane. Half the animals were perfused, and half underwent rapid decapitation. Animals used for immunohistochemistry were perfused with phosphate-buffered saline (PBS) followed by 4% paraformaldehyde. Brains were post-fixed in 4% paraformaldehyde before being switched to 30% sucrose until sectioning. Tissue was sectioned at 30 μm and stored in PBS-azide until staining. Animals used for high performance liquid chromatography (HPLC) underwent rapid decapitation. The hippocampus was dissected, weighed, and stored at-80°C until processing.

### 2.5 High Performance Liquid Chromatography

To verify DSP-4 lesions, HPLC was performed in half of the animals to assess NE tissue concentrations in the hippocampus, as previously described [19,20]. An ESA 5600A CoulArray detection system, coupled with an ESA Model 584 pump and ESA 452 autosampler was used for analysis. Separations were carried out on an MD-150 × 3.2 mm C18 column (3 µm, Thermo Fisher Scientific) maintained at 30°C. The mobile phase, delivered isocratically at 0.4 mL/min, consisted of 8% acetonitrile, 75 mM NaH2PO4, 1.7 mM 1-octanesulfonic acid sodium, and 0.025% trimethylamine, adjusted to pH 2.9. A 20 µL sample was injected, and detection was performed using a 6210 electrochemical cell (ESA, Bedford, MA) equipped with 5020 guard cell. The guard cell potential was set to 475 mV, and the analytical cell potentials were set to−175, 100, 350 and 425 mV. Analytes were identified by comparing their retention times to those of known standards (Sigma Chemical Co., St. Louis MO). Quantification was performed by comparing the peak areas of the analytes to those of standards on the dominant sensor.

### 2.6 Immunohistochemistry

Vendor and species antibody information are listed in Table 1. Hippocampal and prefrontal cortex (PFC) sections were stained with anti-NE transporter (NET) antibody to evaluate LC innervation, 4G8 to measure amyloid-β plaques, and CP13 to label phosphorylated tau, as well as glial fibrillary acidic protein (GFAP) to mark astrocytes and anti-ionized calcium binding adaptor molecule 1 (IBA1) to mark microglia as measures of neuroinflammation.

**Table 1.** Antibody information.

| **Primary Antibody** | **Vendor**<br>**(Catalog #)** | **Dilution** | **Secondary**<br>**Antibody** | **Vendor**<br>**(Catalog #)** | **Dilution** |
| --- | --- | --- | --- | --- | --- |
| Mouse anti-CP13 | Gifted from<br>Peter Davies | 1:2000 | Goat anti-mouse<br>568 | ThermoFisher<br>Scientific<br>(A-11004) | 1:500 |
| Mouse anti-4G8 | BioLegend<br>(800798) | 1:1000 | Goat anti-mouse<br>568 | ThermoFisher<br>Scientific<br>(A-11004) | 1:500 |
| Rabbit anti-IBA1 | Fujifilm<br>(019-1974) | 1:1000 | Goat anti-rabbit<br>488 | ThermoFisher<br>Scientific<br>(A11008) | 1:500 |
| Mouse anti-NET | Mab<br>Technologies<br>(NET05-2) | 1:500 | Goat anti-mouse<br>568 | ThermoFisher<br>Scientific<br>(A-11004) | 1:500 |
| Rabbit anti-GFAP | Abcam<br>(ab7260) | 1:1000 | Goat anti-rabbit<br>488 | ThermoFisher<br>Scientific<br>(A11008) | 1:500 |
| Chicken anti-TH | Abcam<br>(ab76442) | 1:1000 | Goat anti-chicken<br>647 | ThermoFisher<br>Scientific<br>(A-21449) | 1:500 |

Additionally, LC sections were stained with tyrosine hydroxylase (TH) to mark noradrenergic neurons and assess their integrity.

Sections were incubated in blocking solution (5% normal goat serum in 0.1 PBS-Triton) for 1 h. Sections were then incubated in primary antibodies for 24 h at 4°C with a 1:500 (NET), 1:1000 (4G8, IBA1, TH, and GFAP) or 1:2000 (CP13) dilution of primary antibody. Following overnight incubation, sections were washed 3 × 5 min in 1x PBS. They were then covered and incubated in appropriate 1:500 dilution of secondary antibody for 2 h. Sections were subsequently washed for 3 × 5 min in 1x PBS and mounted onto Superfrost Plus slides in weak mounting buffer. After drying, slides were coverslipped with DAPI Fluoromount-G® (Southern Biotech).

### 2.7 Image Analysis

Image analysis was performed in ImageJ/Fiji version 1.54g (National Institutes of Health, USA). Standardized regions of interest (214,200 pixels^2^) across sections in the hippocampus and PFC were used to quantify CP13, IBA1, and GFAP. Images were grayscaled and thresholded using the Otsu method. Amyloid-β plaques were analyzed as a percent area, CP13 as mean gray value, and IBA1 and GFAP as counts using the analyze particles function within subregions of the hippocampus and PFC. For TH analysis in the LC, a standardized ROI was applied, and TH fluorescence was quantified as the mean gray value within the ROI.

Line scan analysis was performed on NET images as previously described[8,10]. A line of standard size was drawn through the image. ImageJ was used to generate line intensity profiles, which were exported to MATLAB (Version R2024b), where intensity data was plotted. MATLAB’s peak detection function was then used to count the number of detected peaks, which corresponded with the number of NET+ fibers.

### 2.8 Statistics and Analysis

Data visualization and statistical analysis were carried out using GraphPad Prism (Version 10.5.0). Except for circadian locomotion and fear conditioning all data were analyzed using a 2-way RM-ANOVA with genotype and treatment as factors. Circadian locomotion and fear conditioning were analyzed using a 3-way RM-ANOVA with time, genotype, and treatment as factors. When interactions were detected, multiple comparisons testing with Šidák correction were performed. A Fisher’s exact test was used in addition to the 2-way ANOVA to analyze novelty-suppressed feeding data.

## 3. Results

### 3.1 DSP-4 reduces hippocampal NE levels and noradrenergic fibers, and damages noradrenergic neurons in the LC

We verified DSP-4 lesions using three independent approaches. First, we performed HPLC on tissue samples taken from the hippocampus (Fig. 1a). There was a main effect of treatment (F_(1, 17)_ = 104.3, p<0.0001), with both WT and TgF344-AD animals that received DSP-4 displaying dramatically decreased concentrations of NE. Second, NET immunoreactivity was used to visualize noradrenergic axons/terminals in the hippocampus and PFC (Fig. 1b–f). There was a main effect of treatment in the PFC (F_(1, 17)_ = 12.24, p = 0.0028) and DG (F_(1, 17)_ = 38.93, p<0.0001), as well as main effects of treatment and genotype in CA1 (Treatment: F_(1, 17)_ = 12.84, p = 0.0023, Genotype: F_(1, 17)_ = 6.462, p = 0.0211) and CA3 (Treatment: F_(1, 17)_ = 14.92, p = 0.0012, Genotype: F_(1, 17)_ = 7.800, p = 0.0125). Across all regions, DSP-4-treated animals had fewer NET+ fibers, and in CA1 and CA3, TgF344-AD rats had reduced NET+ fibers compared to WT littermates. Finally, TH immunoreactivity was used to determine whether our DSP-4 protocol compromised LC somatodendritic integrity (Fig. 1g–h). There was a main effect of treatment (F_(1, 18)_ = 10.01, p = 0.0054), with DSP-4-treated animals displaying reduced TH immunoreactivity in the LC compared to saline-treated animals, regardless of genotype. Taken together, these results indicate that the DSP-4 regimen was effective in triggering LC fiber and cell body/proximal dendrite degeneration.

**Figure 1.**
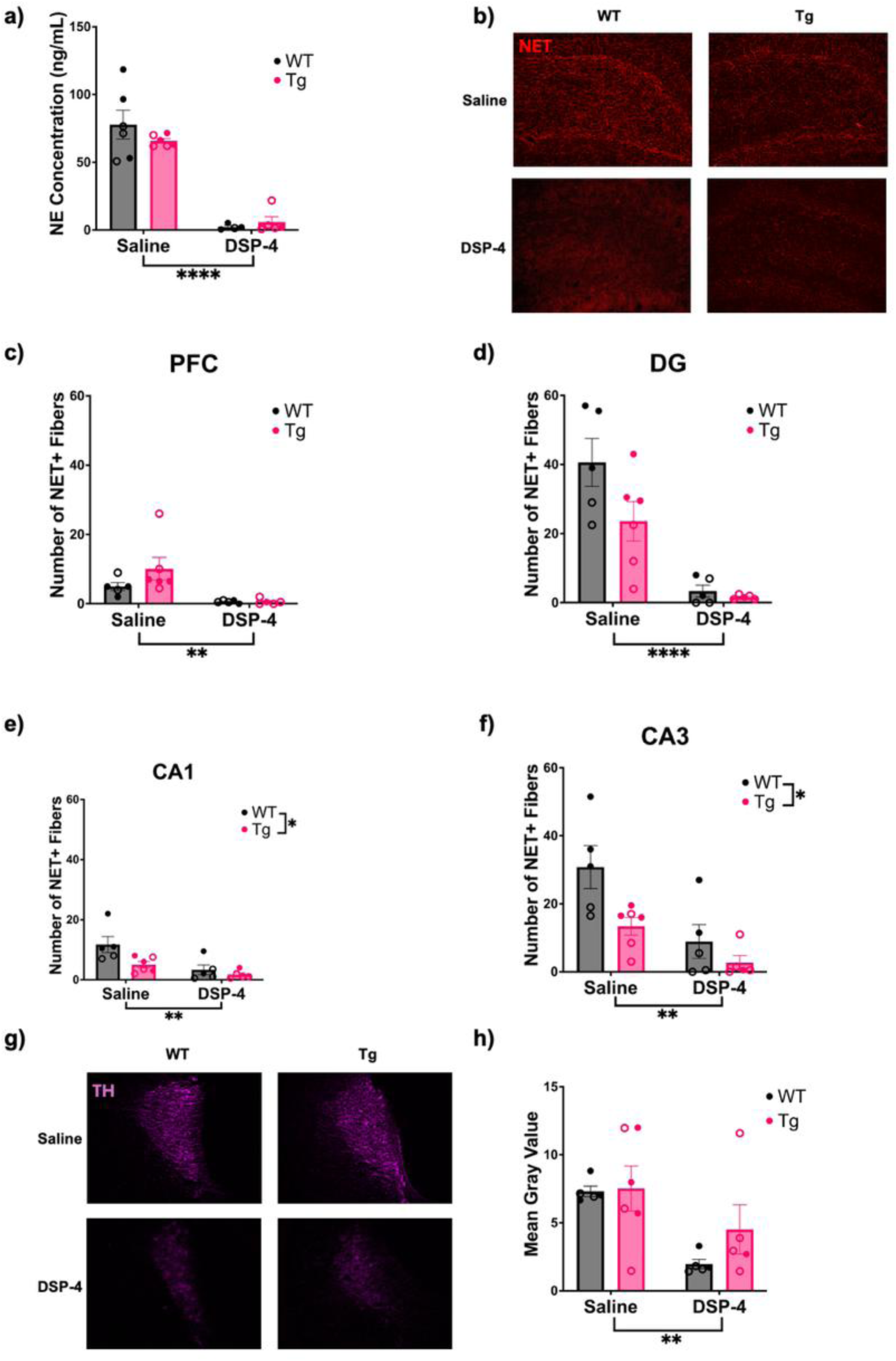
Repeated DSP-4 administration depletes forebrain NE and damages LC cell bodies. WT (black) and TgF344-AD (pink) animals received monthly injections of DSP-4 or saline, and the effects of DSP-4 administration on the LC-NE system were evaluated using HPLC and immunohistochemistry. DSP-4 administration reduced hippocampal NE levels (a). Representative images (40x) of NET+ fibers in the DG (b). Quantification of NET+ fibers in the PFC (c), DG (d), CA1 (e), and CA3 (f). Representative images (40x) of TH staining in the LC (g). Quantification of TH immunoreactivity in the LC expressed as mean gray value (h). Females are represented by open circles and males by closed circles. Data are mean ± SEM; n = 4-6 per group (a) or n = 5-6 per group (c-f, h). **p ≤ 0.01, ****p ≤ 0.0001.

### 3.2 DSP-4 and TgF344-AD genotype each independently affect behavior in a paradigm-specific manner

#### 3.2.1 Social discrimination

A constellation of social deficits presents in not only those with diagnosed AD, but also individuals who go on to develop AD [29,30]. NE also has a broad role in sociability, and more specifically in social memory, where reductions in NE result in social memory deficits [31–33]. We assessed social and non-social preference in a 3-chamber interaction assay, where rats and mice typically display novelty preference. There were no main effects of genotype or treatment on the ratio of time spent investigating the novel object versus novel animal, with all groups preferring interactions with the novel animal (Fig. 2a). There was a main effect of genotype on the ratio of time spent investigating a novel versus familiar animal (F_(1, 39)_ = 8.025, p = 0.0073), with WT animals spending more time investigating the novel animal, while TgF344-AD rats spent more time investigating the familiar animal (Fig. 2b).

**Figure 2.**
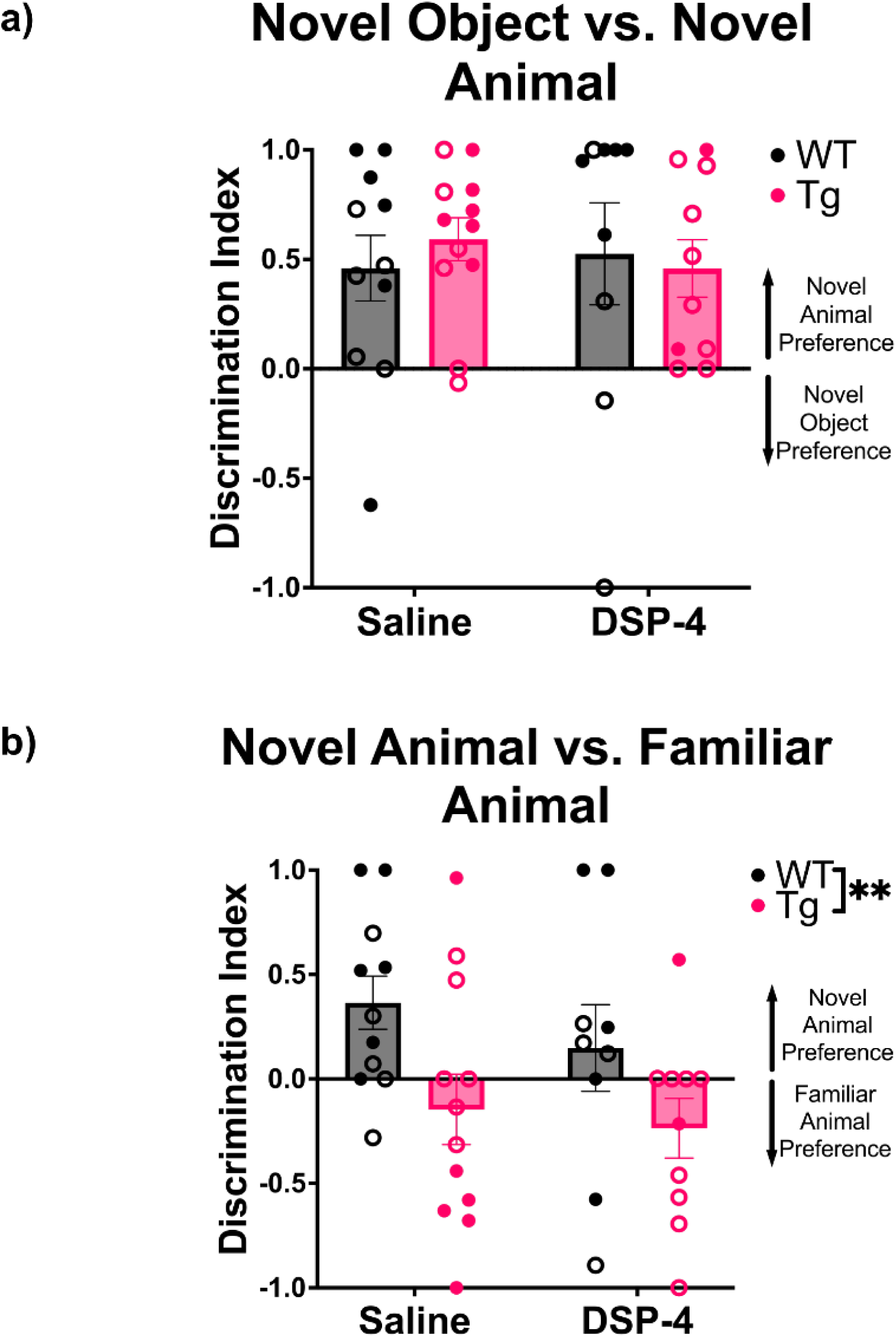
TgF344-AD animals display impaired social discrimination. WT (black) and TgF344-AD (pink) animals received monthly injections of DSP-4 or saline, and the ratio of time spent interacting with a novel object versus a novel animal (a) and a familiar versus novel animal (b) was calculated. All animals displayed a preference for novel animals versus novel objects (a), but only WT animals displayed a preference for novel versus familiar animals (b). Females are represented by open circles and males by closed circles. Data are mean ± SEM; n= 9-12 per group. **p ≤ 0.01.

#### 3.2.2 Circadian locomotor activity

Sleep disturbances are observed during the preclinical and prodromal phases of AD, and there is evidence suggesting a bidirectional relationship between sleep disturbances and AD progression [34,35]. The LC-NE system is a central regulator of arousal and sleep-wake cycles, and has been hypothesized to contribute to sleep disturbances in AD [36–38]. To investigate gross changes in the sleep-wake cycle, we used locomotor activity as a proxy for circadian arousal. There were main effects of time (F_(91, 3496)_ = 10.65, p<0.0001), genotype (F_(1, 3496)_ = 32.21, p<0.0001), and treatment (F_(1, 3496)_ = 12.15, p=0.0005) on locomotor activity (Fig. 3b). In general, WT animals ambulated less than TgF344-AD animals, and DSP-4-treated animals ambulated less than saline-treated animals.

**Figure 3.**
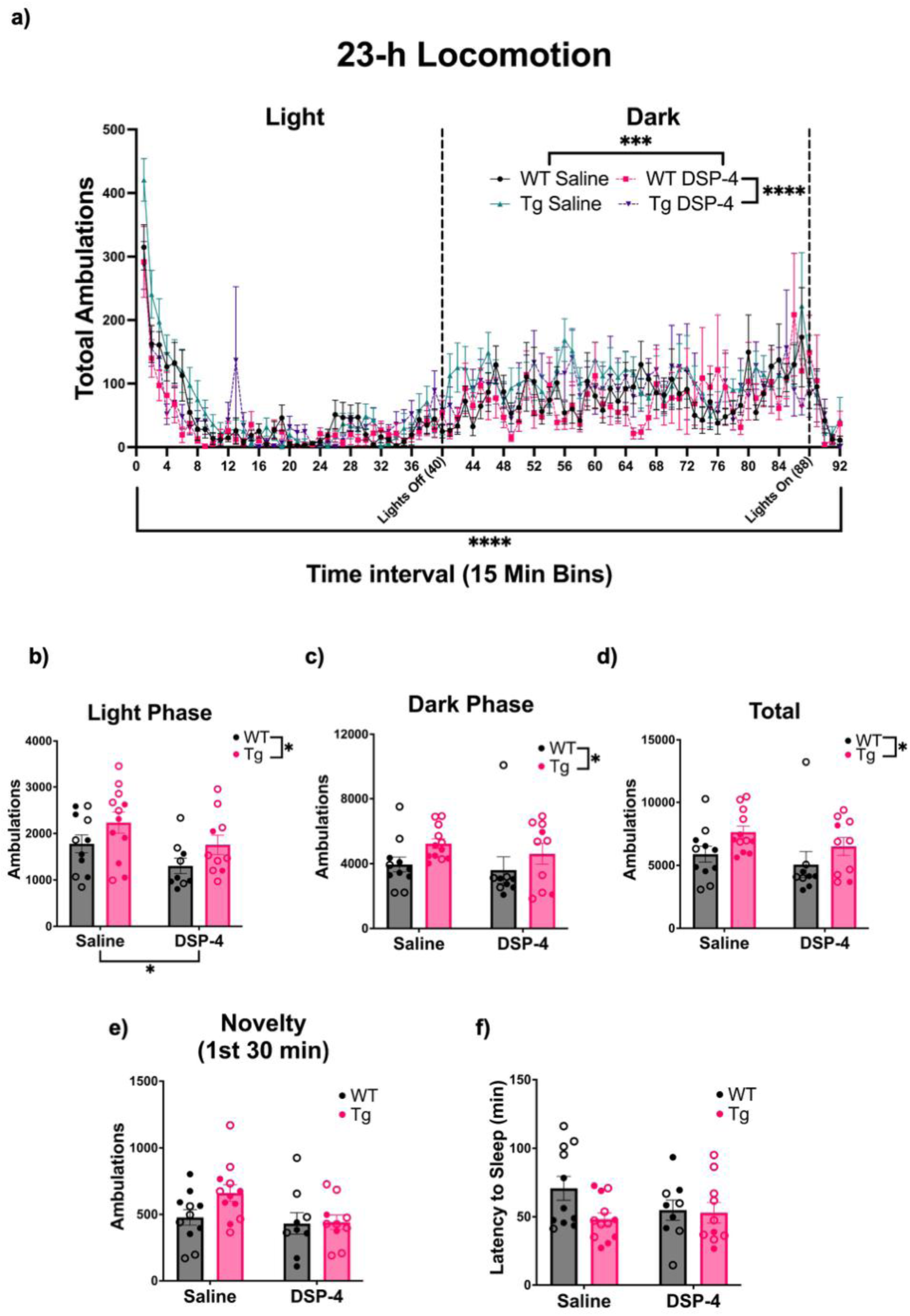
Circadian rhythms and arousal are affected by both genotype and DSP-4 treatment. WT (black) and TgF344-AD (pink) animals received monthly injections of DSP-4 or saline. Circadian activity was evaluated during a 23-h locomotor behavioral paradigm (a). Total ambulations were binned over the light phase (b), dark phase (c), entire 23-h task (d), and first 30 min (e), Arousal was evaluated during sleep latency, where the latency to fall asleep was quantified (f). Females are represented by open circles and males by closed circles. Data are mean ± SEM; n= 9-12 per group. *p ≤ 0.05, ***p ≤ 0.001, ****p ≤ 0.0001.

Locomotion data over the 23-h period was further divided into the light and dark phases (Fig. 3d & c). During the light phase, there were main effects of treatment (F_(1, 38)_ = 5.322, p = 0.0266) and genotype (F_(1, 38)_ = 4.865, p = 0.0335). Saline-treated and TgF344-AD rats were more active than DSP-4-treated and WT animals, respectively. There was also a main effect of genotype (F_(1, 38)_ = 4.294, p=0.0451) during the dark phase, with TgF344-AD animals moving more than WT animals. Over the entire 23-h period, a main effect of genotype (F_(1, 38)_ = 5.097, p = 0.0298) was observed, with TgF344-AD animals showing greater locomotor activity than WT animals (Fig. 3d).

#### 3.2.3 Novelty-induced locomotion and sleep latency

Decreased levels of arousal have been observed in people with mild cognitive impairment, and the LC-NE system plays an important role in regulating arousal [34,38,39]. We used novelty-induced locomotor activity (defined as the first 30 min of the circadian locomotor activity test) and latency to fall asleep following a mild stimulus (gentle handling) to assess arousal. For novelty-induced locomotor activity, there was a main effect of treatment (F_(1, 38)_ = 4.292, p=0.0451), with saline-treated animals ambulating more compared to DSP-4-treated animals, regardless of genotype (Fig. 3e). There were no significant effects of treatment or genotype on latency to fall asleep (Fig. 3f).

#### 3.2.4 Stress-induced repetitive and apathy-like behaviors

Apathy, defined as reduced motivation, activity of daily living, goal-directed behavior, and emotional flattening, is a common prodromal AD symptom, correlates with AD severity, predicts the transition from normal cognition to mild cognitive impairment to AD, and is ameliorated by noradrenergic drugs [35,40,41]. We used stick chewing and nestlet shredding as markers of environmental engagement that rats are typically highly motivated to partake in as proxies for apathy-like behavior. At the same time, the immediate response to cage changes during the start of these tasks can induce stress-related repetitive behaviors that can be assessed simultaneously [26]. For nestlet shredding, there was a main effect of treatment on the percentage of the nestlet shredded when measured after 90 min (F_(1, 38)_ = 7.841, p=0.0080) and 6 days (F_(1, 38)_ = 18.25, p = 0.0001), with saline-treated animals shredding more than DSP-4-treated animals (Fig. 4a & b). Similarly to nestlet shredding, there was a main effect of treatment on stick chewing, with saline-treated animals chewing a greater percentage of the wooden stick than DSP-4-treated animals when measured after 24 h (F_(1, 38)_ = 34.56, p<0.0001) and 1 week (F_(1, 38)_ = 16.90, p=0.0002) (Fig. 4c & d).

**Figure 4.**
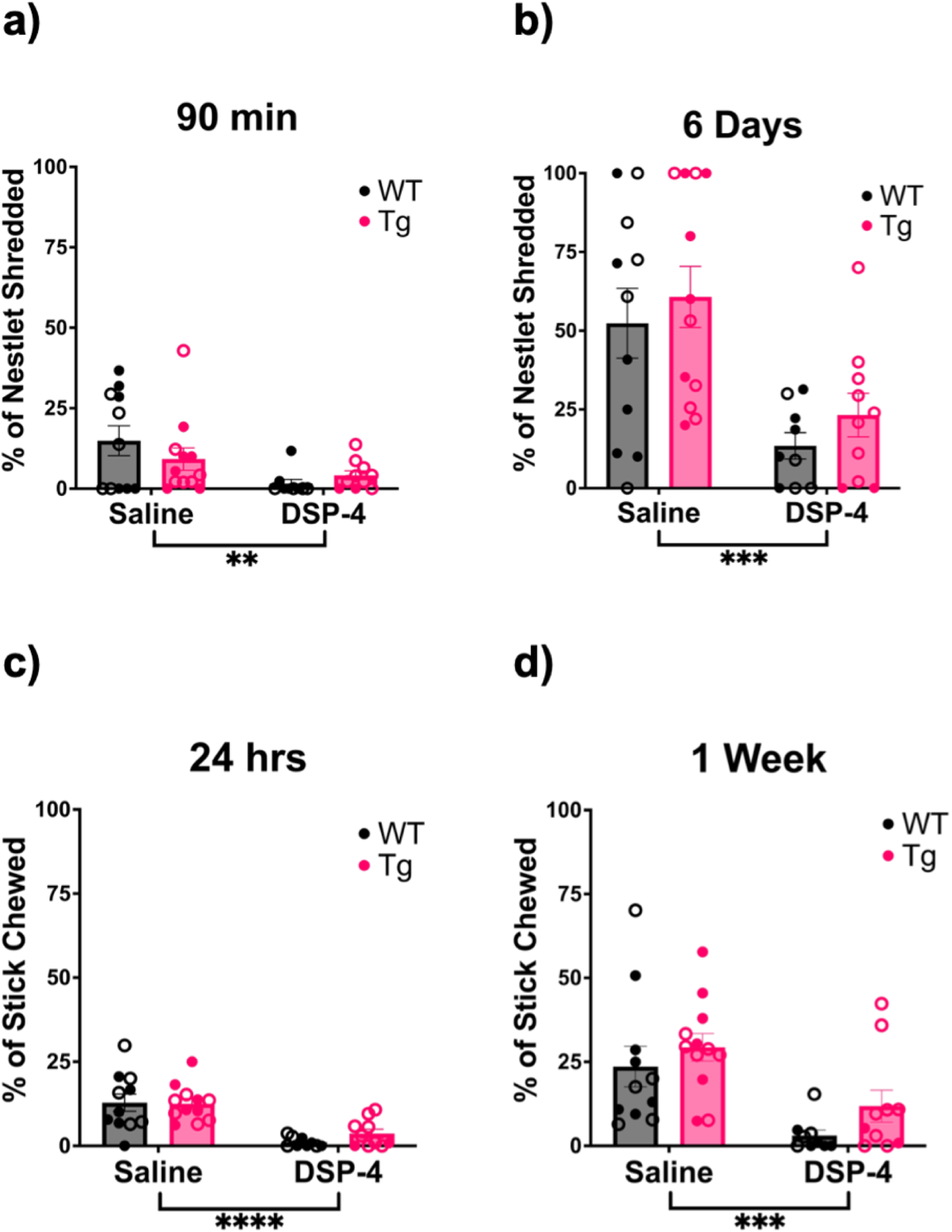
Repeated DSP-4 treatment ablates immediate stress-induced repetitive behavior and causes apathy-like behavior. WT (black) and TgF344-AD (pink) animals received monthly injections of DSP-4 or saline, and stress-induced repetitive and apathy-like behavior was evaluated using nestlet shredding. Quantification of the percentage of nestlet shredded after 90 min (a) and 6 days (b). Apathy-like behavior was also evaluated using a stick chewing behavioral paradigm. Shown are the percentage of stick chewed after 24 hrs (c) and 1 week (d). Animals treated with DSP-4 show a lack of nestlet shredding and stick chewing behavior. Females are represented by open circles and males by closed circles. Data are mean ± SEM; n= 9-12 per group. **p ≤ 0.01, ***p ≤ 0.001, ****p ≤ 0.0001.

#### 3.2.5 Open field and novelty-suppressed feeding

Anxiety is highly prevalent in preclinical AD, and influences how amyloid-β impacts cognitive decline [41–43]. Furthermore, the LC-NE system plays an important role in modulating anxiety [44–46]. We used both the open field and novelty-suppressed feeding tasks to assess anxiety-like phenotypes. These two tasks are distinct conflict-based models of anxiety. The open field test relies on the proclivity of rodents to engage in exploration of novel environments, but also avoid light open spaces. There was a main effect of treatment (F_(1, 36)_ = 4.915, p=0.0330) on the percentage of time spent in the center of the arena during the open field task. DSP-4-treated animals showed increased anxiety-like behavior, spending less time in the center compared to saline-treated controls (Fig. 5a).

**Figure 5.**
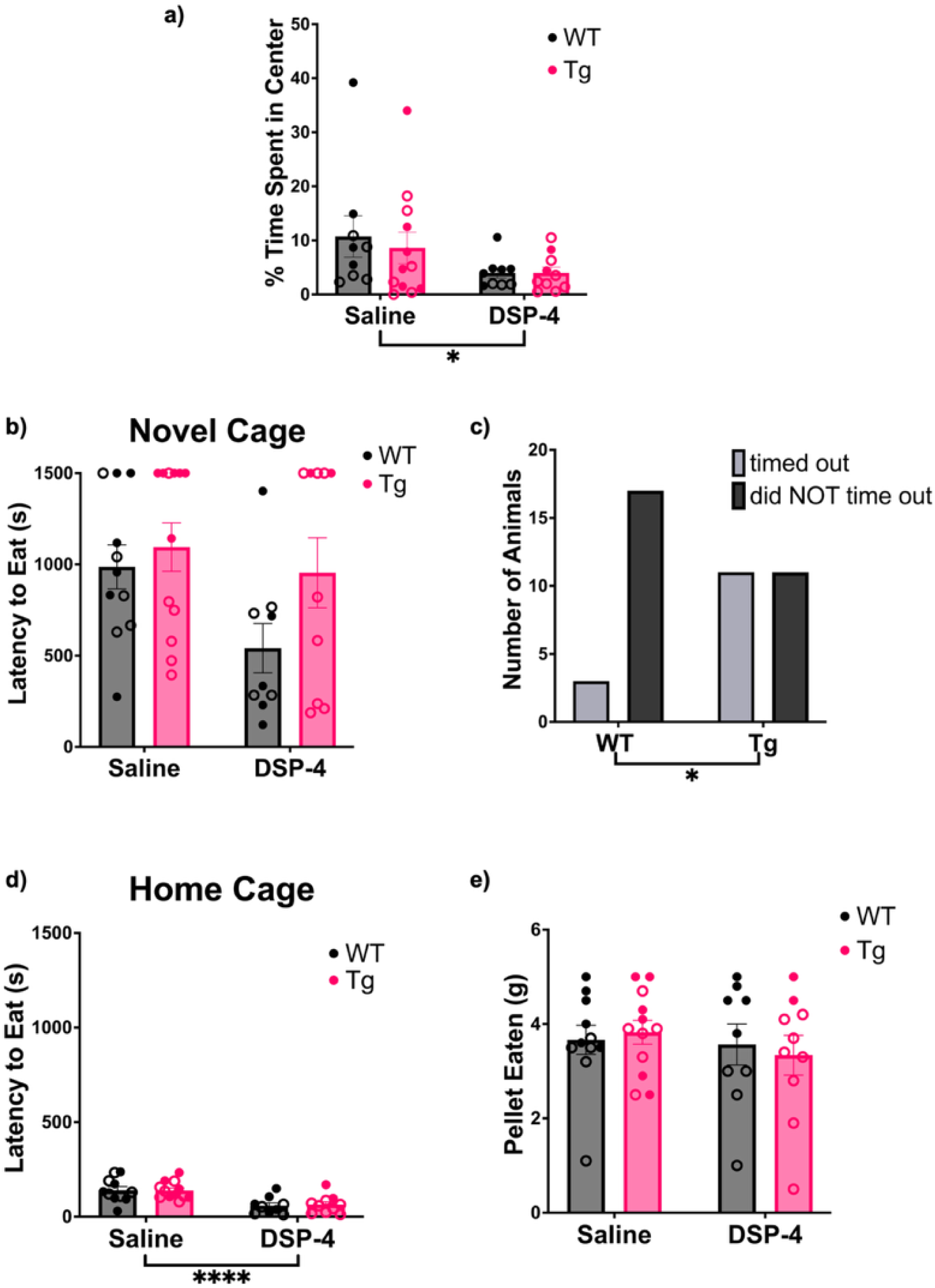
Treatment and genotype cause increased anxiety-like behaviors in a task-dependent manner. WT (black) and TgF344-AD (pink) animals received monthly injections of DSP-4 or saline. Anxiety-like behavior was evaluated using an open field behavioral paradigm where the percentage of time spent in the center of the arena was quantified (a). Anxiety-like behavior was also evaluated during a novelty-suppressed feeding behavioral paradigm. Shown are the latency to eat in the novel cage (b), a comparison of the number of TgF344-AD and WT animals that timed out of the task (c), the latency to eat in the home cage (d), and the total amount of pellet consumed after 1 h (e). Animals treated with DSP-4 showed increased anxiety-like behaviors in the open field task while TgF344-AD animals displayed elevated anxiety-like behavior during novelty-suppressed feeding. Females are represented by open circles and males by closed circles. Data are mean ± SEM; n= 9-12 per group. *p ≤ 0.05, ****p ≤ 0.0001.

Meanwhile, the novelty-suppressed feeding task pits a hungry rat’s motivation to eat against its fear of bright open spaces of unknown safety. We have previously shown that among anxiety tests, the novelty-suppressed feeding task is particularly sensitive to NE perturbations [47]. However, there were no significant main effects or interactions in the latency to eat in the novel environment (Fig. 5b). One caveat to analyzing the novelty-suppressed feeding test is that a time limit can create a ceiling effect. We noted that many animals, especially TgF344-AD rats, timed out during the novelty-suppressed feeding task. To account for this, we re-analyzed the data with a Fisher’s Exact test on the fraction of animals that timed out, and found that significantly more TgF344-AD rats did not bite the food pellet within the allotted time compared to WT controls (p = 0.0232; Fig. 5c). There was also a significant effect of treatment (F_(1, 38)_ = 24.54, p < 0.0001) on the latency to eat in the home cages (Fig. 5d). DSP-4-treated animals ate more quickly compared to the saline-treated group. There was no effect on the total amount of food eaten 1 h after the task (Fig. 5e).

#### 3.2.6 Cue-and context-dependent fear conditioning

Cued and contextual fear conditioning are associative learning tasks that assess distinct cognitive processes. While contextual fear conditioning is hippocampal-dependent, cued fear conditioning is not, and involves other brain regions such as the amygdala [48,49]. AD is characterized by severe memory impairment, including deficits fear conditioning in both patients and animal models [8,25,50]. Importantly, the LC-NE system plays a role in nearly all facets of cued and contextual fear learning [49]. During training, there was a main effect of time (F_(6, 266)_ = 15.58, p < 0.0001), with more freezing evident upon each subsequent shock presentation (Fig. 6a). There were no effects of genotype or treatment during training, suggesting all groups learned the tone-shock association at the same rate. During contextual fear conditioning, there were main effects of time (F_(6, 266)_ = 4.080, p = 0.0006), genotype (F_(1, 266)_ = 12.17, p = 0.0006), and treatment (F_(1, 266)_ = 15.69, p <0.0001; Fig. 6b). There were similar main effects during cued fear conditioning for time (F_(6, 203)_ = 62.44, p < 0.0001), genotype (F_(1, 203)_ = 7.170, p = 0.0080), and treatment (F_(1, 203)_ = 11.94, p = 0.0007; Fig. 6c). In general, TgF344-AD rats and DSP-4-treated rats had increased freezing in both paradigms compared to their WT and saline-treated counterparts, respectively.

**Figure 6.**
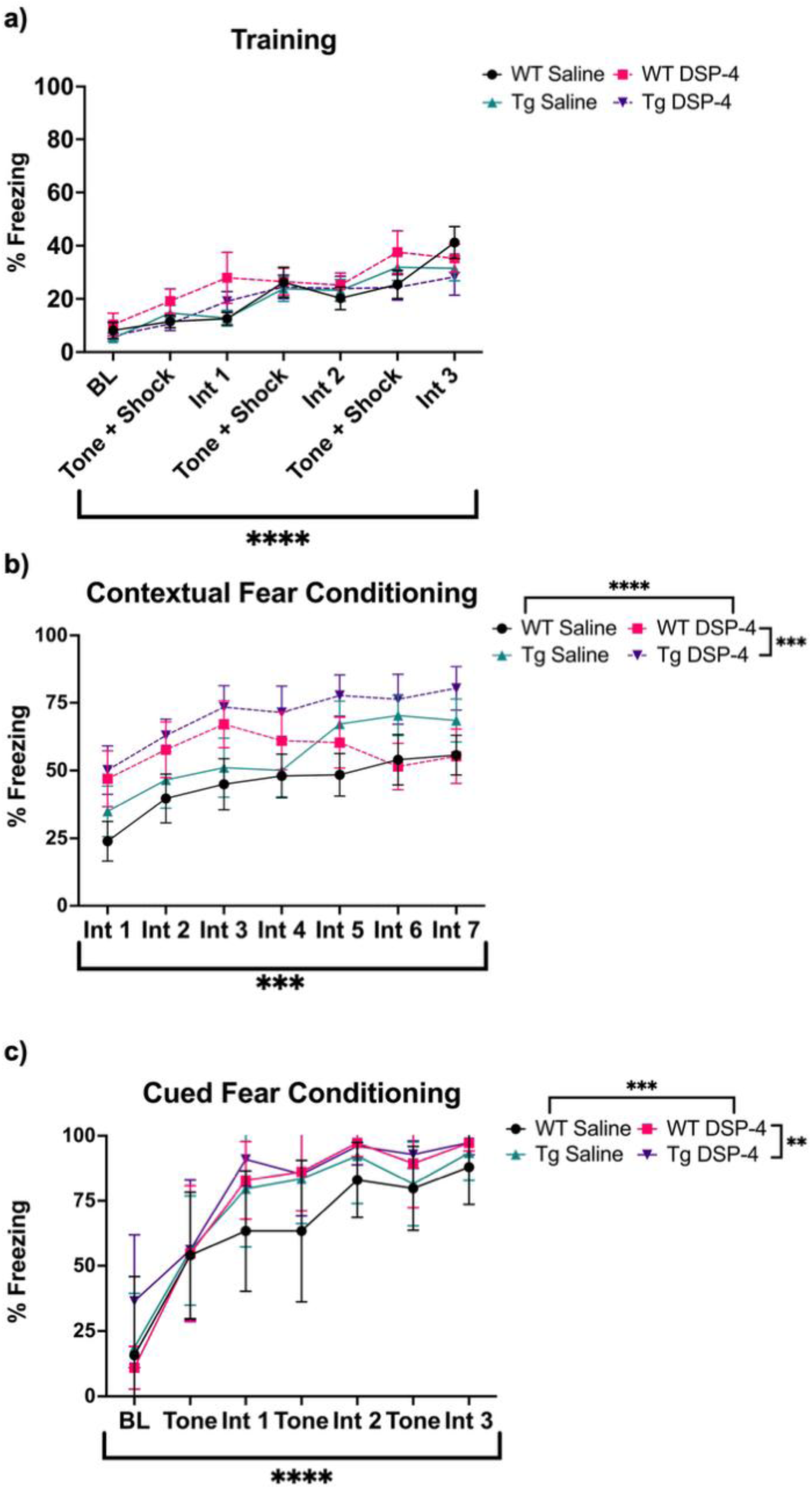
Treatment and genotype increase freezing during contextual and cued fear conditioning. WT (black) and TgF344-AD (pink) animals received monthly injections of DSP-4 or saline, and learning and memory was evaluated using fear conditioning. The percentage of time spent freezing was quantified during training (a), contextual fear conditioning (b), and cued fear conditioning (c). Despite similar rates of associative learning, TgF344-AD rats and animals treated with DSP-4 displayed higher freezing in both contextual and cued fear conditioning. Data are mean ± SEM; n= 9-12 per group. **p ≤ 0.01, ***p ≤ 0.001, ****p ≤ 0.0001.

### 3.3 DSP-4 and TgF344-AD genotype each independently affect pathology in a marker-specific manner

#### 3.3.1 Amyloid

4G8 was used to detect and quantify amyloid-β plaques in the hippocampus and PFC (Fig. 7a–e). There were no significant main effects of genotype, treatment, or their interaction on amount of amyloid pathology (Fig. 7b–e). Overall, very little 4G8 immunoreactivity was observed, but when present was detected only in TgF344-AD animals, consistent with prior reports using similarly aged animals [8,10,25,51,52]. DSP-4 treatment did not exacerbate amyloid-β pathology.

**Figure 7.**
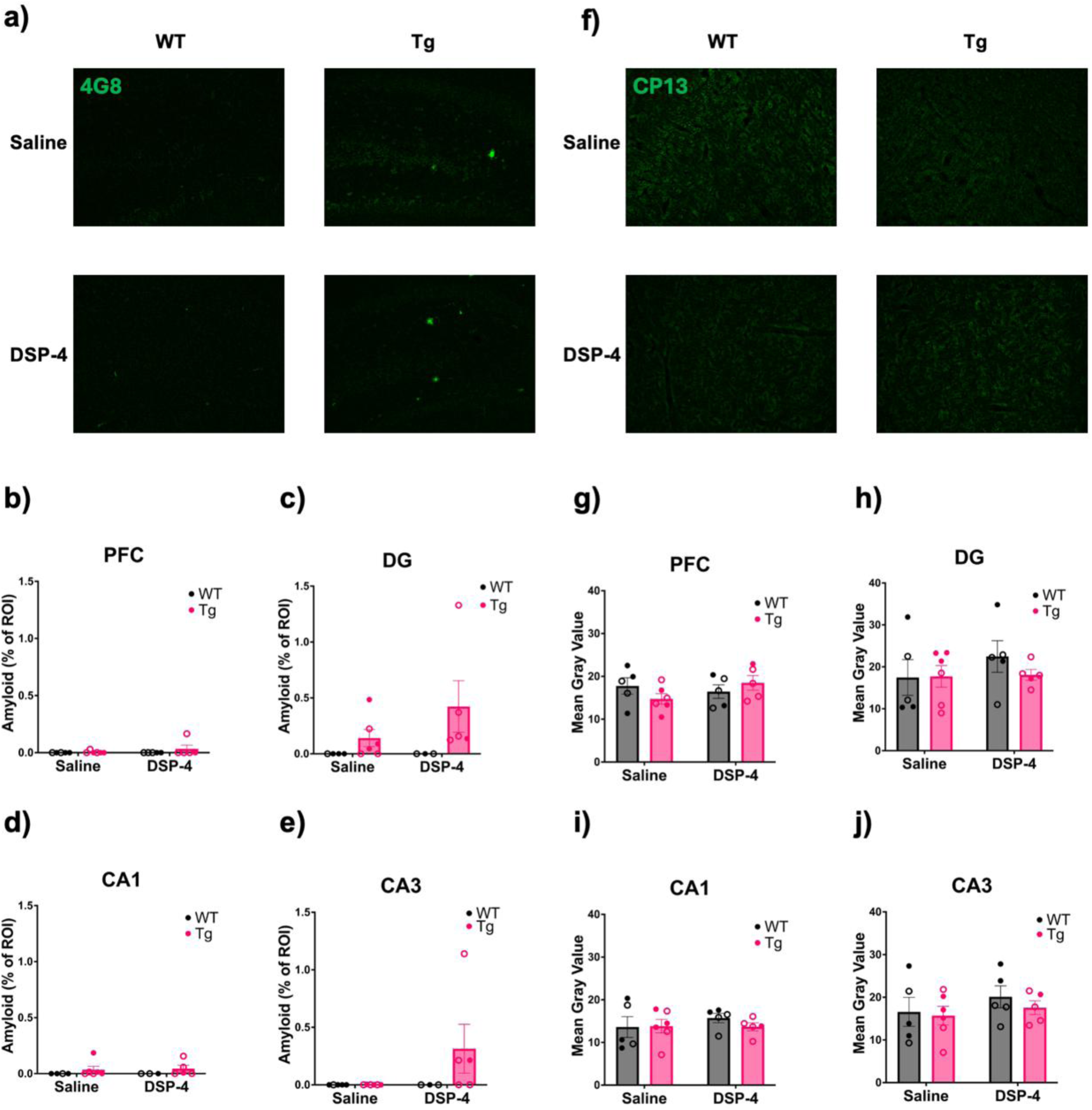
Neither treatment nor genotype impacted AD-related pathology. WT (black) and TgF344-AD (pink) animals received monthly injections of DSP-4 or saline, and amyloid-β and hyperphosphorylated tau pathology was quantified. Representative images (40x) of amyloid-β pathology in the DG (a). Quantification of amyloid-β pathology in the PFC (b), DG (c), CA1 (d), and CA3 (e). Representative images (40x) of CP13 staining in the PFC (f). Quantification of hyperphosphorylated tau pathology in the PFC (g), DG (h), CA1 (i), and CA3 (j) expressed as the mean gray value within a standardized ROI. Neither treatment nor genotype exacerbated AD-related neuropathology. Females are represented by open circles and males by closed circles. Data are mean ± SEM.

#### 3.3.2 Hyperphosphorylated tau

We next used the CP13 Ser202 phospho-antibody to stain and quantify hyperphosphorylated tau in the hippocampus and PFC (Fig. 7f–j). CP13 immunoreactivity in TgF344-AD rats was sparse at this age, in agreement with previous results from our lab using similarly aged animals [8,25]. Similar to amyloid-β pathology, DSP-4 treatment did not worsen tau pathology, and there were no significant main effects or interactions.

#### 3.3.3 Microglia

IBA1 immunoreactivity was used to assess microglial neuroinflammation in the hippocampus and PFC (Fig. 8a–e). There was a main effect of genotype in the PFC (F_(1, 17)_ = 5.795, p = 0.0277; Fig. 8b) and CA1 (F_(1, 15)_ = 5.680, p = 0.0308; Fig. 8d), with WT animals displaying higher microglia counts compared to TgF344-AD animals. No significant main effects or interactions were observed in the DG (Fig. 8c) or CA3 (Fig. 8e).

**Figure 8.**
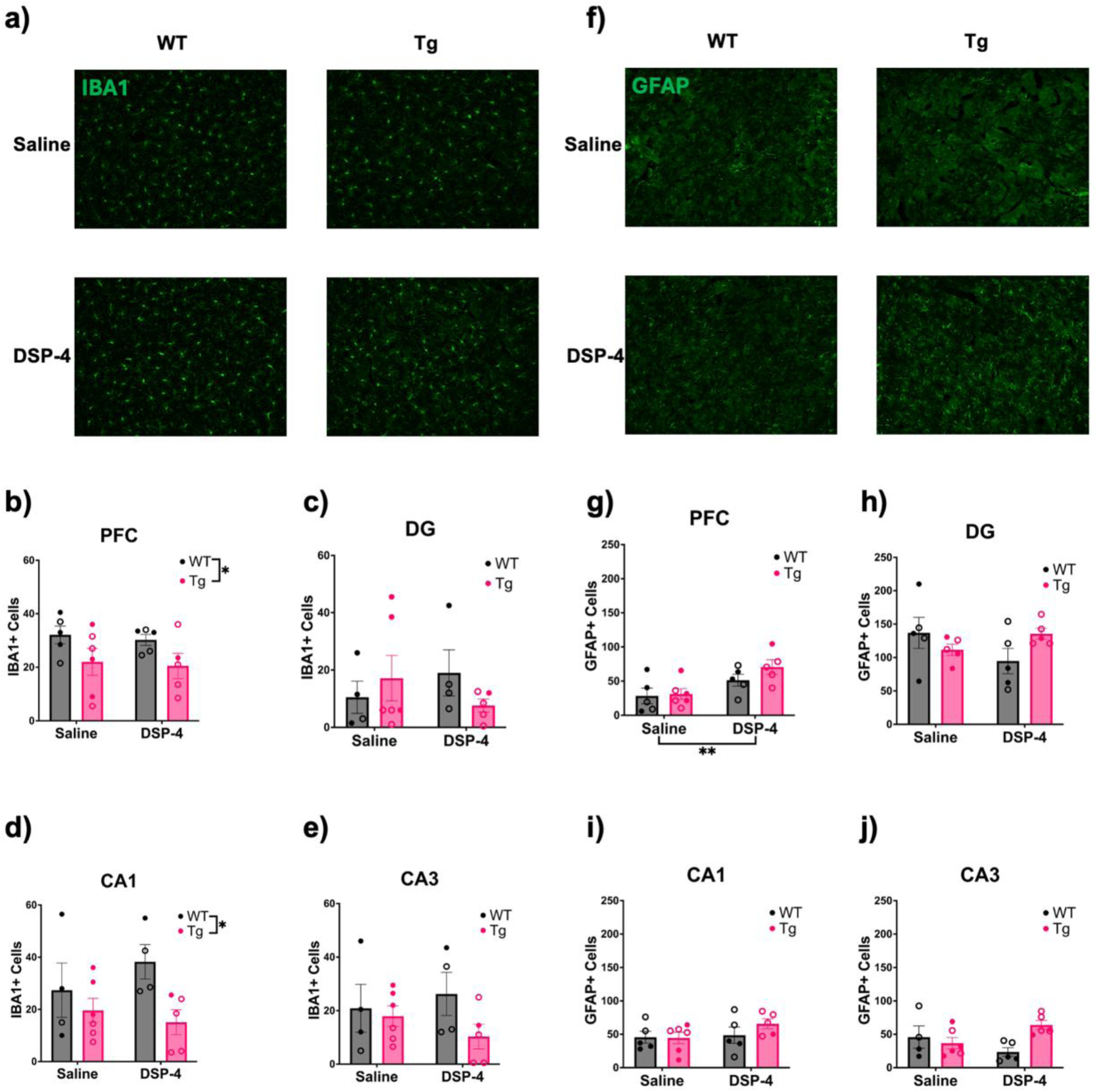
Levels of neuroinflammation depend on treatment and genotype. WT (black) and TgF344-AD (pink) animals received monthly injections of DSP-4 or saline. IBA1 and GFAP immunoreactivity was evaluated. Shown are representative images (40x) of IBA1 staining in the PFC (a). Quantification of IBA1 in the PFC (b), DG (c), CA1 (d), and CA3 (e). Representative images (40x) of GFAP staining in the PFC (f). Quantification of GFAP in the PFC (g), DG (h), CA1 (i), and CA3 (j). WT rats displayed more IBA1+ cells compared to TgF344-AD rats in the PFC and CA1 while DSP-4 treatment caused increased GFAP+ cells in the PFC. Females are represented by open circles and males by closed circles. Data are mean ± SEM; n= 5-6 per group. *p ≤ 0.05, **p ≤ 0.01.

#### 3.3.4 Astrocytes

GFAP was used to stain for astrocytic neuroinflammation (Fig. 8f–j). There was a main effect of treatment in the PFC (F_(1, 17)_ = 10.60, p=0.0046; Fig. 8g), with DSP-4-treated rats showing higher GFAP immunoreactivity than their saline-treated counterparts. There were no significant main effects or interactions in the DG (Fig. 8h) or CA1 (Fig. 8i). There was a significant genotype x treatment interaction in CA3 (F_(1, 16)_ = 6.658, p = 0.0201; Fig. 8j), where it appeared that DSP-4 reduced GFAP immunoreactivity in WT animals and increased it in TgF344-AD animals, but Šidák-corrected post-hoc analyses did not reveal any significant pairwise differences.

## 4. Discussion

We investigated the consequences of repeated DSP-4 administration during pre-pathological stages on molecular and behavioral phenotypes in TgF344-AD rats. We validated DSP-4 lesions by showing that our paradigm reduced tissue NE levels, lowered NET+ fiber innervation, and decreased TH immunofluorescence in the LC. We expected lower LC-NE integrity would interact with TgF344-AD genotype to exacerbate molecular and behavioral phenotypes.

However, although both DSP-4 treatment and TgF344-AD genotype independently influenced behavior and molecular pathology in a paradigm-and marker-specific manner, almost no genotype x treatment interactions were observed.

The social behavior paradigm revealed a deficit in TgF344-AD rats. While both WT and TgF344-AD rats preferred investigating novel animals over novel objects, TgF344-AD rats preferred investigating the familiar animal over the novel one, suggesting a deficit in social memory and consistent with clinical features of AD and results from other rodent AD models [53–55]. Our results are also aligned with recent studies reporting similar deficits across social tasks in TgF344-AD rats between 6-14 months of age [56,57], indicating that social deficits are a prominent early non-cognitive feature of AD. DSP-4 did not impact social behavior. One report in APP23 transgenic mice showed social discrimination deficits only following DSP-4 treatment, suggesting an interaction between pathology and NE denervation on social deficits [16].

We next assessed circadian rhythms and arousal using 23-h locomotor activity and sleep latency. TgF344-AD animals displayed increased locomotor activity compared to WT animals, similar to mice with the same transgenes [53]. However, our prior study with 6-month-old TgF344-AD rats did not reveal locomotor hyperactivity; instead, 12-month WT and TgF344-AD rats showed reduced locomotion, suggesting an age-dependence to the phenotype [8]. This age dependence is supported by another report, though these effects also depended on sex [51].

These complex results necessitate a well-controlled study to understand the interaction between sex, age, and genotype that obscures clear interpretation of our findings. Meanwhile, DSP-4 reduced locomotion, which agrees with our previous report [19] but contrasts with other evidence indicating no effect of DSP-4 on locomotor activity [20].

The lack of effect during sleep latency was surprising, given the role that NE plays in modulating the sleep/wake cycle and the prevalence of sleep disturbances in AD. However, we reported a similar failure of TgF344-AD genotype to influence sleep latency [8]. To our knowledge, this is the first study using sleep latency to probe the impact of DSP-4 on arousal, and we observed no differences resulting from LC lesions. Quantifying aspects of non-REM and REM with EEG, as well as examining sleep over a longer epoch that includes both light and dark cycles, may reveal differences not detectable at the level of this behavioral assay [58].

We also uncovered an effect of DSP-4 on stress-induced repetitive behavior, with DSP-4-treated animals shredding significantly less than saline-treated animals. We previously reported that stress-induced repetitive behaviors are attenuated in NE-deficient (*Dbh* −/−) mice, which are restored with pharmacological augmentation of NE [26]. Although repetitive behaviors are highly prevalent in AD [59–61], genotype did not affect stress-induced shredding. Many repetitive behaviors displayed by AD patients are verbal [59,61], indicating that nestlet shredding may not be appropriate for assessing repetitive behaviors. Instead, we propose monitoring ultrasonic vocalizations, which may be more stereotyped in TgF344-AD rats and could be linked to LC-NE dysfunction.

We extended the nestlet shredding test to 6 days and up to a week with stick chewing to assess environmental engagement to proxy apathy-like behavior, and found that DSP-4-treated animals did not shred or chew as much as saline-treated animals. Given the role of the LC-NE system in environmental engagement and motivation, the emergence of apathy-like behaviors following DSP-4 lesions is unsurprising [62]. However, the lack of phenotype in TgF344-AD rats contrasts with another report in similarly aged rats, suggesting a task-specific nature to these behaviors [63]. We did not score nest building behaviors, which could have unveiled changes in health and well-being [64].

We found increased anxiety-like behavior in TgF344-AD rats in the novelty-suppressed feeding task, consistent with our prior report [8]. While there were no significant effects or interactions in the latency to eat in a novel environment, significantly more TgF344-AD rats failed to eat within the allotted time, confirming increased anxiety-like behavior.

DSP-4-treated animals displayed higher anxiety-like behavior during the open field task. This contrasts with previous work showing that DSP-4 does not alter open field behavior in WT or AD mice [19,20]. However, our results are in line with prior rat work showing that DSP-4 increases anxiety-like behavior in an open field-like setting [65].

DSP-4 also caused a trend towards decreased latency to eat in the novelty-suppressed feeding task, particularly in WT rats. This is similar to *Dbh-/-* mice, which display a shorter latency to eat in a novel environment [47], but contrasts with previous results indicating that DSP-4 increases latency to eat during this task in mice [19]. The latter result may be due to the different DSP-4 treatment regimens employed compared to our study. There was also an unexpected, robust effect of DSP-4 causing shorter latency to eat in the home cage in the absence of changes in overall hunger levels assessed by total amount eaten within an hour. Given the complex role of the LC-NE system in feeding [66,67], more work needs to be done to understand the effects of LC lesions in this task. However, based on our current and prior studies, the novelty-suppressed feeding task appears particularly sensitive to changes in NE transmission [8,26,66].

Typically, young TgF344-AD rats do not display fear conditioning deficits, a measure of associative learning[8,52]. Because LC degeneration is correlated with cognitive deficits in AD, we hypothesized that DSP-4 may unmask this phenotype [13,68]. However, TgF344-AD and DSP-4-treated animals froze *more* during both cued and contextual fear conditioning in the absence of differences in acquisition. This could result from LC-NE hyperactivity or may reflect increased anxiety-like behavior, as shown in other assays. For example, 6-month TgF344-AD rats exhibit elevated spontaneous bursting and evoked LC activity [9]. These changes at the level of the cell bodies coincide with downstream β-AR hypersensitivity [52], which is associated with maintenance of hippocampal-dependent memory. Moreover, a single dose of DSP-4 can increase downstream α2-AR binding sites up to a month after administration, and a similar mechanism could be at play with β-ARs [18].

We hypothesized that DSP-4 would exacerbate neuroinflammation, amyloid-β, and hyperphosphorylated tau [15–17,20]. While we observed some evidence of increased neuroinflammation due to DSP-4, there were no significant changes in AD pathology.

Small amounts of 4G8 immunoreactivity were present in TgF344-AD animals, consistent with the low levels of amyloid-β pathology observed at 6 months [8,10,25,51,52]. There was a non-significant trend towards increased amyloid-β burden in DSP-4-treated animals. There was also no significant difference in CP13 staining. This is unsurprising, as forebrain hyperphosphorylated tau levels are typically low in TgF344-AD rats until older ages [10,25].

Microglia and astrocyte activation are key components of neuroinflammation in AD [69–71].

Reduced NE signaling is necessary for microglia to increase their process extension and surveillance [69,72]. NE levels decline in AD, potentially resulting in a loss of suppression that could lead to excessive neuroinflammation [8,10,14,69–72]. Consistent with this hypothesis, DSP-4-treated animals displayed higher astrocyte counts in the PFC. There was no significant effect of DSP-4 on microglia counts in the hippocampus. Instead, we found a significant effect of genotype, with WT animals exhibiting significantly greater microglia counts in the PFC and CA1, consistent with our prior report [8].

Most previous studies using AD models examined the effects of LC-NE degeneration in much older animals, which exacerbated pre-existing AD-like neuropathology and behavioral deficits [16,20,22–24]. Instead, we examined TgF344-AD rats at ∼5-6 months when there is little AD-like neuropathology in the forebrain. When considered with prior studies, our data suggests that LC-NE lesions exacerbate behavioral and molecular phenotypes in later stages of disease when pathology is already present. Model choice may also affect outcomes, as DSP-4 accelerated AD-related phenotypes in McGill-R-Thy1-APP transgenic rats [15]. Another report using a single stereotaxic injection of DβH-saporin to lesion the LC in similarly aged TgF344-AD rats showed accelerated cognitive decline and pathology, but the magnitude of these changes compared to WT conditions is unknown[73].

Considering our findings in the context of other DSP-4 studies, it is important to consider that treatment paradigms can impact extent of damage [15–21]. Furthermore, while DSP-4 administration reduced TH immunoreactivity, we did not conduct cell counts and thus cannot conclude whether LC cells were degenerating or merely lost their molecular identity [19]. A loss of proximal dendrites also could have contributed to the lower TH immunoreactivity in the LC, which could also explain the reduction of LC volume but not number of cells in the earliest stages of clinical AD [6]. Finally, while DSP-4 is considered LC-specific[74], other evidence refutes this notion [18], and the contribution of dysfunction of noradrenergic nuclei other than the LC (e.g. A1, A2) to AD-related behavioral and molecular phenotypes is underexplored.

Our study expanded the behavioral and molecular characterization of TgF344-AD rats in response to early LC-NE lesions. TgF344-AD genotype and DSP-4 rarely interacted to exacerbate AD-related symptoms or pathology, having greater effects when considered independently. Further characterization in older TgF344-AD rats and using different DSP-4 administration paradigms are needed to untangle the relationship between LC lesions and AD progression.

## Acknowledgements

This work was supported by the Emory HPLC Bioanalytical Core (RRID:SCR_023531) and the Emory University Rodent Behavioral Core (RRID:SCR_012499).

## Author Contributions

Alexia Marriott (Conceptualization; Data curation; Formal analysis; Investigation; Methodology; Software; Validation; Visualization; Writing – original draft; Writing – review & editing); Jason Schroeder (Investigation; Writing – Review and Editing); Anu Korukonda (Investigation; Writing – Review and Editing); Brittany Pate (Investigation; Writing – Review and Editing) Katharine McCann (Investigation; Writing – Review and Editing); David Weinshenker (Conceptualization; Formal analysis; Funding acquisition; Methodology; Resources; Supervision; Validation; Visualization; Writing – Original Draft; Writing – Review and Editing); Michael Kelberman (Conceptualization; Data curation; Formal analysis; Funding acquisition; Investigation; Methodology; Resources; Software; Supervision; Validation; Writing – original draft; Writing – Review and Editing)

## Ethical Considerations

The Emory University Institutional Animal Care and Use Committee approved the experimental procedures used in this study (PROTO201700315 - NE and Neurodegeneration).

## Consent to Participate

Not applicable

## Consent for Publication

Not applicable

## Declaration of Conflicting Interests

The author(s) declared no potential conflicts of interest with respect to the research, authorship, and/or publication of this article.

## Funding

This work was supported by funding from the National Institute on Aging (AG062581 to DW, AG069502 to MAK, and AG066511 to MAK), National Institute of Neurological Disorders and Stroke (NS96050 to MAK).

## Data availability

All data will be made available upon request to the corresponding author.

## Notes

### Competing Interest Statement

The authors have declared no competing interest.

